# Irgm2 and Gate-16 cooperatively dampen targeting of caspase-11 to Gram-negative bacterial products

**DOI:** 10.1101/2020.03.06.980433

**Authors:** Elif Eren, Rémi Planès, Salimata Bagayoko, Pierre-Jean Bordignon, Karima Chaoui, Audrey Hessel, Karin Santoni, Miriam Pinilla, Odile Burlet-Schiltz, Jonathan C. Howard, Masahiro Yamamoto, Etienne Meunier

## Abstract

Inflammatory caspase-11 (rodent) and caspases-4 and -5 (human) detect gram-negative bacterial component LPS in the host cell cytosol, which promotes activation of the non-canonical inflammasome. Although non-canonical inflammasome-induced pyroptosis and IL-1 related cytokine release is of importance to mount an efficient immune response against various bacteria, its unrestrained activation drives sepsis. This suggests that cellular components might tightly control the threshold level of the non-canonical inflammasome in order to ensure efficient but not deleterious inflammatory response. Here we show that the IFN-inducible protein Irgm2 and the ATG8 family member Gate-16 cooperatively slow down non-canonical inflammasome activation both in macrophages and *in vivo*. Specifically, the Irgm2/Gate-16 axis dampens caspase-11 targeting to intracellular bacteria, which lower caspase-11-mediated pyroptosis and cytokine release. Specifically, deficiency in *Irgm2* or *Gate16* opens an alternative road for caspase-11 targeting to intracellular bacteria, independently of the classical pathway driven by the Guanylate Binding Proteins (GBPs). Thus, our findings provide new molecular effectors involved at fine-tuning the optimal non-canonical inflammasome response and add novel insights in the understanding of the immune pathways they control.

## Introduction

Inflammasomes are cytosolic innate immune complexes that initiate inflammatory responses upon sensing of microbe- and damage-associated molecular patterns (MAMPs and DAMPs respectively) (Hayward *et al*, 2018). Specifically, the rodent caspase-11 (and its human orthologous Caspases-4 and -5) detects the presence of the gram-negative bacterial cell wall component lipopolysacharide (LPS) in the host cell cytosol (Hagar *et al*, 2013; Kayagaki *et al*, 2013, 2011; Yang *et al*, 2015; Aachoui *et al*, 2013; Broz *et al*, 2012). LPS interaction with caspase-11 caspase activation and recruitment domains (CARD) domain promotes its oligomerization and activation, which triggers the activation of the non-canonical inflammasome (Yang *et al*, 2015). Upon activation (Lee *et al*, 2018), caspase-11 cleaves and activate the pyroptosis executioner gasderminD (gsdmD) into a p30 active fragment (Shi *et al*, 2015; Kayagaki *et al*, 2015). Cleaved gsdmD then forms a pore into PIP2-enriched domains at the plasma membrane, which triggers pyroptosis, a pro-inflammatory form of cell death (Sborgi *et al*, 2016; Aglietti *et al*, 2016; Liu *et al*, 2016; Shi *et al*, 2015). In parallel, gsdmD pores-induced ionic perturbations also trigger activation of the canonical NLRP3 inflammasome, which ultimates in the caspase1-dependent maturation of the pro-inflammatory cytokines IL-1β/IL-18 (Schmid-Burgk *et al*, 2015; Rühl & Broz, 2015; Kayagaki *et al*, 2011). Although caspase-11 confers host protection against intracellular Gram-negative bacteria (Cerqueira *et al*, 2018; Chen *et al*, 2018; Aachoui *et al*, 2013), unrestrained caspase-11 response drives host pathology that includes irreversible organ failure, blood clothing and sepsis (Yang *et al*, 2019; Cheng *et al*, 2017; Napier *et al*, 2016; Deng *et al*, 2018; Yang *et al*, 2019; Kayagaki *et al*, 2011, 2013, 2015; Rathinam *et al*, 2019). This suggests that host regulators might fine-tune the non-canonical inflammasome in order to optimize caspase11-dependent response. So far, only few of them were described including SERPINB1-inhibited caspase-11/-4 /-1 activation in resting cells or ESCRT-mediated plasma membrane repair (Choi *et al*, 2019; Rühl *et al*, 2018).

Crucial at regulating the activation the non-canonical inflammasome pathway are the IFN-inducible GTPases, the so-called Guanylate Binding Proteins (GBPs) and the Interferon Regulated GTPase (Irg) Irgb10 (Lagrange *et al*, 2018; Man *et al*, 2016, 2015, Meunier *et al*, 2014, 2015; Pilla *et al*, 2014; Finethy *et al*, 2015; Zwack *et al*, 2017; Wallet *et al*, 2017; Costa Franco *et al*, 2018; Liu *et al*, 2018; Cerqueira *et al*, 2018). Specifically, GBPs (1, 2 and 5) are recruited on LPS-enriched structures such as cytosolic gram-negative bacteria and their derived products Outer Membrane Vesicles (OMVs) (Meunier *et al*, 2014; Finethy *et al*, 2017; Fisch *et al*, 2019; Santos *et al*, 2018; Lagrange *et al*, 2018; Man *et al*, 2015). Recruited GBPs then enrol caspase-11 that will bind the LPS moiety Lipid A, hence promoting the non-canonical inflammasome pathway (Fisch *et al*, 2019). Beyond their role at triggering GBP expression, IFNs induce more than 2000 antimicrobial genes (Green *et al*, 2018). Among them, many IFN-inducible regulatory genes also counter-balance over activation of the cells (Green *et al*, 2018). For instance, SOCS1 and USP18 are ISGs that balance the level of the host cell response (Basters *et al*, 2018; Liau *et al*, 2018). In this context, we hypothesized that IFNs, in addition to their ability to promote GBP expression, might also induce negative regulators of the non-canonical inflammasome. In this regard, Irgm proteins belong to the IFN-inducible Interferon-Related GTPases (Irgs) family proteins (Kim *et al*, 2012; Pilla-Moffett *et al*, 2016; Kim *et al*, 2019). Human express one IRGM protein, with various spliced variants, that is not IFN-inducible but that requires IFN signalling to be functional (Kim *et al*, 2019). By contrast, mice express three different Irgms, namely Irgm1, 2 and 3 (Kim *et al*, 2012). All Irgms lack the ability to hydrolyse the GTP due to a mutation in their catalytic domain (GMS) whereas other Irgs are GTPase active (GKS) (Coers, 2013). Although complementary processes occur, previous studies underscored an inhibitory role of Irgm1 and 3 on the recruitment and/or activation of the GBPs and Irg-GKS on microbial membranes (Haldar *et al*, 2015; Feeley *et al*, 2017; Haldar *et al*, 2013). In addition, recent studies identified Irgm1 and its human homologous IRGM, as being critical for the NLRP3 canonical inflammasome regulation by modulating the autophagy pathway, suggesting a close link between Irgm proteins and inflammasomes (pei *et al*, 2017; Mehto *et al*, 2019a, 2019b). In this context, we hypothesized that Irgms proteins might be IFN-inducible controllers of the non-canonical inflammasome activation threshold.

Here, we report that IFN-inducible Irgm2 and the non-canonical autophagy effector Gate-16 cooperate to slow down non-canonical inflammasome activation by inhibiting caspase-11 targeting to intracellular bacteria.

## Results

### The IFN-inducible protein Irgm2 specifically refrains non-canonical inflammasome response

IFN-inducible Irgms control Irg and GBP microbicidal activity against intracellular pathogens (Pilla-Moffett *et al*, 2016). In this context, we sought to determine whether Irgms might also modulate the non-canonical inflammasome response. Using an RNA interference approach (siRNA), we silenced the three murine Irgms in primary murine bone-marrow derived macrophages (BMDMs) and measured their ability to undergo into caspase-11-dependent cell death and IL-1β maturation upon *Salmonella* Typhimurium challenge. To ensure that the inflammasome response in macrophages is caspase-11-dependent, we used an isogenic mutant of *Salmonella* (*orgA*^-^) lacking expression of SP1 encoded T3SS secretion system (Broz *et al*, 2012). As previously published, *Casp11* and *Gbp2* silencing reduced macrophage death and IL-1β release after 16 hours of infection (**Fig 1A, EV 1A**) (Meunier *et al*, 2014). Importantly, *Irgm2*-silenced BMDMs died and released IL-1β in higher rates than control macrophages (**Fig 1A, EV 1A**). Such process was specific to Irgm2 as *Irgm1* and *3*-targeted siRNA did not induce significant variation in macrophage death and IL-1β release upon *Salmonella* (*orgA*^-^) infection although mRNA levels were efficiently reduced (**Fig 1A, EV 1A**). To further validate that Irgm2 is a regulator of the non-canonical inflammasome response, we challenged WT, *Irgm2*^−/−^, *Casp11*^−/−^ and *GBP*^Chr3–/–^ BMDMs with a panel of gram-negative bacteria that all activate the non-canonical inflammasome. Immunoblotting experiments in WT and *Irgm2*^−/−^ BMDMs showed that Irgm2 is IFN-inducible and that *Irgm2* deficiency does not lead to a defect in caspase-1, -11, GBP2 or GBP5 expression, all involved in the non-canonical inflammasome pathway (**EV1B, C**). Yet, when challenged with various gram-negative bacteria, *Irgm2*^−/−^ macrophages showed an exacerbated cell death, IL-1β release and gasdermin D p30/caspase-1 p20 active fragments compared to their WT counterparts **(Figs 1B, C)**. In addition, Irgm2-regulated cell pyroptosis upon gram-negative bacterial treatment was independent of NLRP3 as the use of the NLRP3 inhibitor MCC950 or *Nlrp3*^−/−^ BMDMs did not drive any defect in cell death (LDH release) while significantly reduced NLRP3-dependent IL-1β release **(EV 1D)**. As expected, both *Casp11*^−/−^ and *GBP*^Chr3– /–^ BMDMs were protected against gram-negative bacteria-induced non-canonical inflammasome response **(Figs 1B, C)**. Importantly, CRISPR-deleted *Irgm2* gene expression in immortalized (i) *Casp11*^−/−^ BMDMs (referred as *Casp11*^−/−^sg*Irgm2*) did not reinduce pyroptosis and IL-1β release upon gram-negative bacterial infections (*S*. Typhimurium *orgA*^-^ and *E. coli*) or *E. coli*-derived outer membrane vesicles (OMVs) exposure, thus confirming that Irgm2 negatively regulated caspase-11-dependent response (**Fig 1D**).

**Figure 1:**
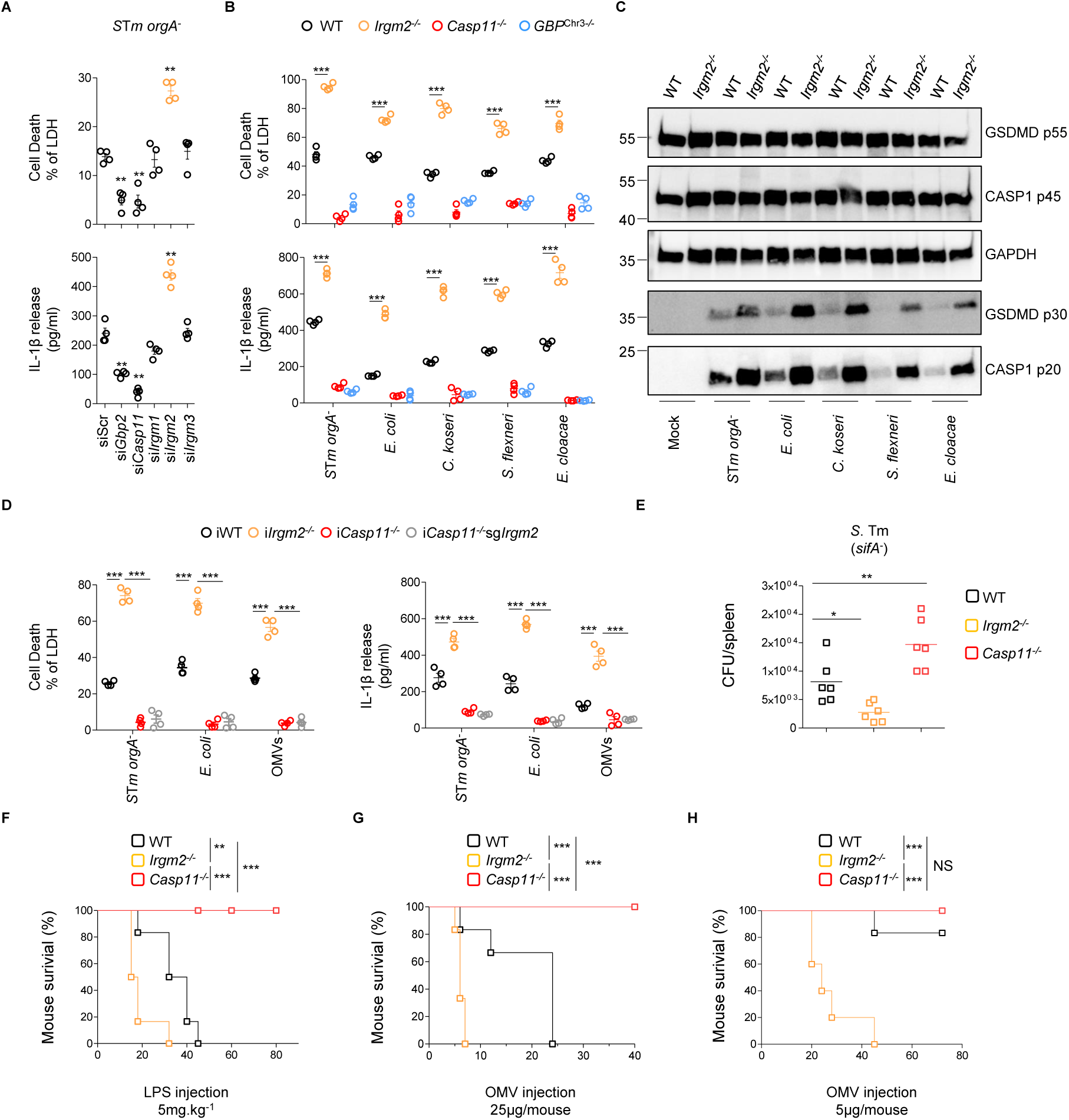
The IFN-inducible protein Irgm2 refrains non-canonical but not canonical inflammasome response. Otherwise specified, BMDMs were either infected with various gram-negative bacterial strains (MOI25) or stimulated with outer membrane vesicles (OMVs) for 16 hours. **(A)** siRNA-treated BMDMs were infected for 16 hours with *S*. Typhimurium (*orgA*^-^) and LDH and IL-1β release were measured. **(B)** Cell death (LDH) and IL-1β release evaluation in WT, *Irgm2*^−/−^, *GBP*^Chr3–/–^ and *Casp11*^−/−^ BMDMs infected for 16 hours with different gram-negative bacteria (MOI 25). **(C)** Western blot examination of processed caspase-1 (p20) and gasdermin-D (p30) in supernatants and pro-caspase-1 (p45), pro-gasdermin-D (p55) and GAPDH in cell lysates of WT and *Irgm2*^−/−^ BMDMs infected for 16 h with different gram-negative bacterial strains. * Non specific **(D)** IL-1β and cell death (% LDH) evaluation immortalized (i) BMDMs, WT, *Irgm2*^−/−^, *Casp11*^−/−^ and *Casp11*^−/−^*Irgm2*^−/−^ (referred as sgIrgm2) after 16 hours of *E. coli, S*. Typhimurium *orgA*^-^ and OMV treatment. **(E)** Spleen bacterial loads from WT (n=6 mice), *Casp11*^−/−^ (n= 6 mice) and *Irgm2*^−/−^ (n= 5 mice) intraperitoneally infected with 3.10^5^ *S*.Tm *sifA*^-^ for 48 hours. **(F-H)** Survival of WT, Casp11^−/−^, *Irgm2*^−/−^ and *GBP*^Chr3–/–^ mice primed with 100µg poly(I:C) for 6 hours and injected (i.p.) with 5mg/k-1 LPS or 5 and 25 µg of OMVs (n=6 animals per condition). Data information: Data shown as means ± SEM (Graphs A, B and D) from n=4 independent pooled experiments; *p ≤ 0.05; **p ≤ 0.01, ***p ≤ 0.001 for the indicated comparisons using t-test with bonferroni correction. Image (C) is representative of one experiment performed three times. (E-H) are representative of three independent experiments; *p ≤ 0.05; **p ≤ 0.01, ***p ≤ 0.001, Mann-Whitney analysis test (E) and log-rank Cox-Mantel test for survival comparisons (F-H).

Then, we sought to determine whether Irgm2 also inhibited canonical inflammasomes. We treated WT, *Irgm2*^−/−^, *Casp11*^−/−^ and *Casp1*^−/−^*Casp11*^−/−^ BMDMs with various inflammasome activators, including flagellin (NLRC4), poly dAdT (AIM2), Nigericin and TcdB (NLRP3), and measured their ability to go into pyroptosis and to release IL-1β. Although all canonical inflammasome activators induced significant caspase-1-dependent response, cell death and IL-1β release levels remained similar in both WT and *Irgm2*^−/−^ BMDMs (**EV 1F**). In addition, activation of the NLRC4 inflammasome by T3SS-expressing *P. aeruginosa* and *S*. Typhimurium remained similar between WT and *Irgm2*^−/−^ BMDMs (**EV 1G**), suggesting that Irgm2 specifically regulates activation of the non-canonical inflammasome.

Caspase-11 is of importance at controlling intracellular gram-negative bacterial infections. Therefore, we performed a previously published model of mouse infection using the cytosolic prone strain *S*. Typhimurium lacking the virulence effector *sifA*^-^ and evaluated the bacterial loads in WT, *Casp11*^−/−^ and *Irgm2*^−/−^ spleens (**Fig 1E**) (Chen *et al*, 2018). We observed increased bacterial loads in spleens from *Casp11*^−/−^ mice compared to WT mice, which is reminiscent of the findings of Chen *et* al., (Chen *et al*, 2018). Yet, spleens from *Irgm2*^−/−^ mice had lower amount of *Salmonella*, thus suggesting that increased non-canonical inflammasome response in *Irgm2*^−/−^ mice was protective against *S*. Typhimurium *sifA*^-^ infection. As Caspase-11 also drives mouse susceptibility to LPS-induced damages, we evaluated whether *Irgm2* deficiency might also sensitize mice to sepsis. We used two LPS-dependent sepsis models, where WT, *Irgm2*^−/−^ and *Casp11*^−/−^ mice were intraperitonally injected with poly(IC) to induce ISG expression (Kayagaki *et al*, 2013; Santos *et al*, 2018). Then, mice were injected either with pure LPS (5mg/kg) or with OMVs (25 µg/mL) (Santos *et al*, 2018; Vanaja *et al*, 2016). Mouse survival showed that while *Casp11*^−/−^ mice had resistance to LPS- and OMV-induced sepsis, WT mice did succumb faster, hence validating or sepsis model (**Fig 1F, G**). We noticed that *Irgm2*^−/−^ mice were even more susceptible than WT mice to both LPS- and OMV-induced sepsis (**Fig 1F, G**). Therefore, we used a sub-lethal model of OMV-induced sepsis by injecting 5µg of OMVs in mice. In such model, both WT and *Casp11*^−/−^ mice recovered from OMV injection whereas all *Irgm2*^−/−^ mice did succumb within 40h post injection (**Fig 1H**). In addition, cytokines assays showed that *Irgm2*^−/−^ mice had an exacerbated release of all pro-inflammatory and inflammasome-related cytokines tested upon OMV challenge, a phenotype that was reduced in *Casp11*^−/−^ mice, hence confirming that Irgm2 expression is crucial to temperate the activation level of the non-canonical inflammasome (**EV 1H**). Altogether, our data suggest that the IFN-inducible Irgm2 specifically refrains the caspase-11 non-canonical inflammasome pathway, which protects mice to LPS-induced sepsis.

### Irgm2 deficiency drives GBP-independent caspase-11 targeting to gram-negative bacterial membranes

IFN-inducible GBPs are important regulators of the non-canonical inflammasome response. Specifically, GBPs-1 and -2 regulate human Caspase -4/-5 activation while GBPs-2 and -5 control mouse caspase 11. Therefore, we hypothesized that Irgm2 might control caspase-11 response through the modulation of the GBPs. Hence, we silenced *Irgm2* in WT and *GBP*^Chr3–/–^ BMDMs (lacking 5 GBPs, 1-3, 5 and 7) and evaluated the caspase-11 response upon OMV stimulation (**EV 2A)**. While OMV-induced both cell death and IL-1β release was strongly reduced in *GBP*^Chr3–/–^, *Irgm2*-silenced *GBP*^Chr3–/–^ BMDMs partially recovered a caspase-11-dependent response, suggesting that Irgm2-inhibited caspase-11 response is independent of GBP modulation (**Fig 2A**). Other and we previously showed that GBPs also controlled canonical AIM2 inflammasome activation upon *F. tularensis* spp *novicida* infection. In this context, we evaluated the importance of Irgm2 at controlling AIM2 inflammasome response upon *F. novicida* infection. Surprisingly, IL-1β and cell death levels were not different between WT and *Irgm2*^−/−^, although they were strongly reduced in *Casp1*^−/−^ *Casp11*^−/−^ and *GBP*^Chr3–/–^ macrophages (**Fig 2B, EV 2B**). In addition, we observed that *Irgm2*-silenced *GBP*^Chr3–/–^ BMDMs did not recover an inflammasome response upon *F. novicida* infection. Then, we generated i*GBP*^Chr3–/–^*Irgm2*^−/−^ (referred as i*GBP*^Chr3–/–^ sg*Irgm2*) by crispr Cas9 and evaluated their response upon *S*. Tm (*orgA*^-^) challenge. i*Irgm2*^−/−^ BMDMs showed time-dependent increased cell death compared to iWT cells (**EV 2C, D**). While, cell death in i*GBP*^Chr3–/–^ BMDMs was strongly impaired, it was partially reversed in i*GBP*^Chr3–/–^sg*Irgm2*, hence suggesting that *Irgm2* deficiency was sufficient to specifically promote caspase-11-dependent response in absence of GBPs (**EV 2C, D**). GBP enrichment on microbial ligand is of importance for efficient caspase-11 and human caspase-4 recruitment (Thurston *et al*, 2016; Fisch *et al*, 2019). However, monitoring for GBP loading on mCherry-expressing *S*. Typhimurim did not show a significant change in the percentage of bacteria targeted by GBP2 (10-15%) in WT and *Irgm2*^−/−^ BMDMs (**Fig 2C**).

**Figure 2:**
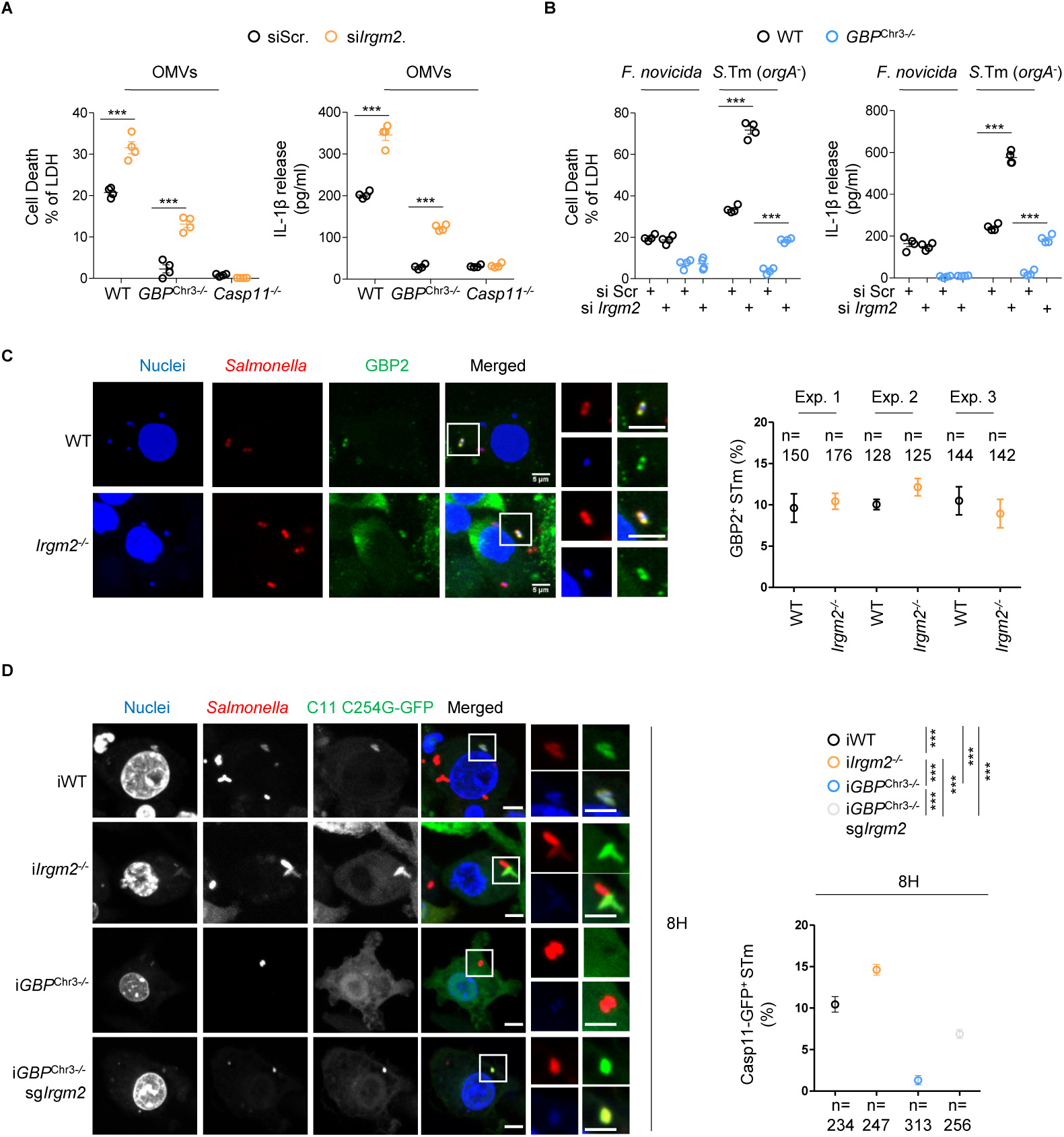
*Irgm2* deficiency opens a GBP-independent road for caspase-11 recruitment to intracellular bacteria. Otherwise specified, BMDMs were treated with 2,5 µg/2.10^5^ cells of OMVs or infected with either *S*. Typhimurium *orgA*^-^ (*S*.Tm *orgA*^-^) or *F. tularensis* spp *novicida* (*F. novicida*) with an MOI of 25 for various times. **(A)** Measure of LDH and IL-1β release in WT, *Irgm2*^−/−^, *GBP*^Chr3–/–^ and *Casp11*^−/−^ BMDMs were Irgm2 was knocked down 16 hours after exposure to 2,5 µg/2.10^5^ cells of OMVs. **(B)** Cell death (LDH) and IL-1β release evaluation in *Irgm2*-silenced WT and *GBP*^Chr3– /–^ BMDMs infected for 16 hours with either *S*.Tm *orgA*^-^ or *F. novicida* (MOI 25). **(C)** Florescence microscopy and associated quantifications of GBP-2 and -5 (green) recruitments to intracellular *S*. Tm *orgA*^-^-mCherry (MOI 10, red) in IFNγ-primed WT, *Irgm2*^−/−^and *Casp11*^−/−^ BMDMs. Nucleus was stained with Hoechst (blue). Confocal images shown are from one experiment and are representative of n=3 independent experiments; scale bars 5 µm. For quantifications, the percentage of GBP-associated bacteria was quantified; quantifications from n=3 independent experiments were then ploted and expressed as mean ± SEM. ***p ≤ 0.001 for the indicated comparisons using t-test with bonferroni correction. **(D)** Confocal fluorescence microscopy images and associated quantifications of caspase-11-C254G-GFP (green) recruitment to *S*.Tm-mCherry (*orgA*^-^, red) in IFNγ-primed iWT, i*Irgm2*^−/−^ i*GBP*^Chr3–/–^ and i*GBP*^Chr3–/–^*Irgm2*^−/−^ BMDMs after 8 hours of infection. Nucleus (blue) was stained with Hoescht; scale bar 5µm. For quantifications, the percentage of bacteria positive for caspase-11-C254G-GFP was determined by combining the bacterial counts from n=3 independent experiments and expressed as mean ± SEM. ***p ≤ 0.001 for the indicated comparisons using t-test with bonferroni correction. Data information: Data shown as means ± SEM (Graphs A, B) from n=4 independent pooled experiments; *p ≤ 0.05; **p ≤ 0.01, ***p ≤ 0.001 for the indicated comparisons using t-test with bonferroni correction. Images (C, D) are representative of one experiment performed three times.

As Caspase-11 activation needs binding to LPS, we hypothesized that Irgm2 might regulate caspase-11 recruitment to bacterial LPS. In order to monitor this, we generated WT, *Irgm2*^−/−^, *GBP*^Chr3–/–^ and *GBP*^Chr3–/–^*Irgm2*^−/−^ iBMDMs that expressed a catalytically inactive mutant of caspase-11 coupled to GFP (Thurston *et al*, 2016) and primed them with IFNγ to induce ISG expression. Observation of CASP11-C254G-GFP recruitment on *S.* Typhimurium (*orgA*^-^) occurred after 4 hours of infection in i*Irgm2*^−/−^, whereas the percentage of caspase-11-targeted bacteria in iWT *GBP*^Chr3–/–^ and *GBP*^Chr3–/–^*Irgm2*^−/−^ iBMDMs remained low or null (**EV 2E**). Yet, after 8 hours of infection, the levels of CASP11-C254G-GFP-associated bacteria increased in iWT (10%), albeit the percentage of CASP11-C254G-GFP^+^ bacteria remained below those observed in *Irgm2*^−/−^ cells (15-16%) (**Fig. 2D**). Strikingly, we noticed that CASP11-C254G-GFP targeting to *Salmonella* was partially restored in *GBP*^Chr3–/–^*Irgm2*^−/−^ iBMDM after 8 hours of infection, although it was strongly impaired in *GBP*^Chr3–/–^ cells (**Fig 2D**). Altogether, our results point out that Irgm2 deficiency opens an alternative road to the GBPs for caspase-11 recruitment on bacterial membranes.

### Irgm2 cooperates with GATE16 to dampen the non-canonical inflammasome activation

As *Irgm2* deficiency increased caspase-11 activation independently of GBPs, we next wondered about additional regulators. We used a GFP-Trap coupled to mass spectrometry (MS) strategy using IFNγ-primed i*Irgm2*^−/−^ BMDMs complemented with a doxycycline-inducible Irgm2-GFP construct (**Fig 3A**). We noticed that one protein was reproductively enriched in the Irgm2-GFP fraction, namely gamma-aminobutyric acid (GABA)-A-receptor-associated protein (GabarapL2 or Gate-16) (**Fig 3A, EV 3A**). Co-immunoprecipitation experiments confirmed that GabarapL2 was enriched in the Irgm2-GFP fraction, hence validating our MS results (**Fig 3B**). Gabarap proteins (Gabarap, GabarapL1 and L2) belong to the ATG8 superfamily proteins, all involved in autophagy/membrane remodelling regulation. While *Gabarap* deficiency leads to increased canonical NLRP3 inflammasome response in mice, there is no information regarding the putative function of Gate-16 at regulating the non-canonical inflammasome. In this context, we found that silencing of *gabarapL2* (*Gate-16*), but not other *gabaraps*, increased OMV-induced caspase-11-dependent cell death and IL-1β release (**Fig 3C, EV 3B, C**). As a control, *Gate-16* silencing did not alter BMDM response to canonical NLRP3 inflammasome activators (**EV 3D**). Then, we sought to determine whether Gate-16-inhibited caspase-11 response was part of the Irgm2 path. Consequently, we silenced *Gate-16* gene expression in WT, *GBP*^Chr3–/–^, *Casp11*^−/−^ and *Irgm2*^−/−^ BMDMs and evaluated the ability of OMVs to induce a caspase-11-dependent response. *Gate-16* silencing in WT BMDMs increased the non-canonical inflammasome response while *Casp11*^−/−^ macrophages remained unresponsive to OMV-induced cell death, IL-1β release (**Fig 3D**). Interestingly, we observed that *GBP*^Chr3–/–^ macrophages silenced for Gate-16 partially recovered their ability to respond to caspase-11 activators (**Fig 3E**). Finally, *Gate-16* knock down in *Irgm2*^−/−^ BMDMs did not exacerbate cell death and IL-1β release nor gasdermin-D or caspase-1 cleavages (**Fig 3D, E**), suggesting that Irgm2 and Gate-16 work together to refrain non-canonical inflammasome response. As *Irgm2* deficiency drives hyper caspase-11 recruitment to bacterial membranes, we wondered about the role of Gate-16 in this process. We silenced *Gate-16* in iWT-expressing CASP11-C254G-GFP and checked for its recruitment on *S*. Tm membranes. Consequently, iWT BMDMs knocked down for *Gate-16* had a more pronounced accumulation of caspase-11 on *S*. Tm membranes than the controls after 4 and 8 hours of infection (**Fig 3F**), which mirrored what we previously observed in *Irgm2*^−/−^ macrophages. To the contrary, Gate16 silencing in *Irgm2*^−/−^ did not increase the percentage of bacteria targeted by CASP11-C254G-GFP after 4 hours of infection (**EV 3E**). Altogether, these results suggest that Irgm2 and Gate-16 cooperate to restrict the non-canonical inflammasome response by refraining caspase-11 targeting to bacterial membranes.

**Figure 3:**
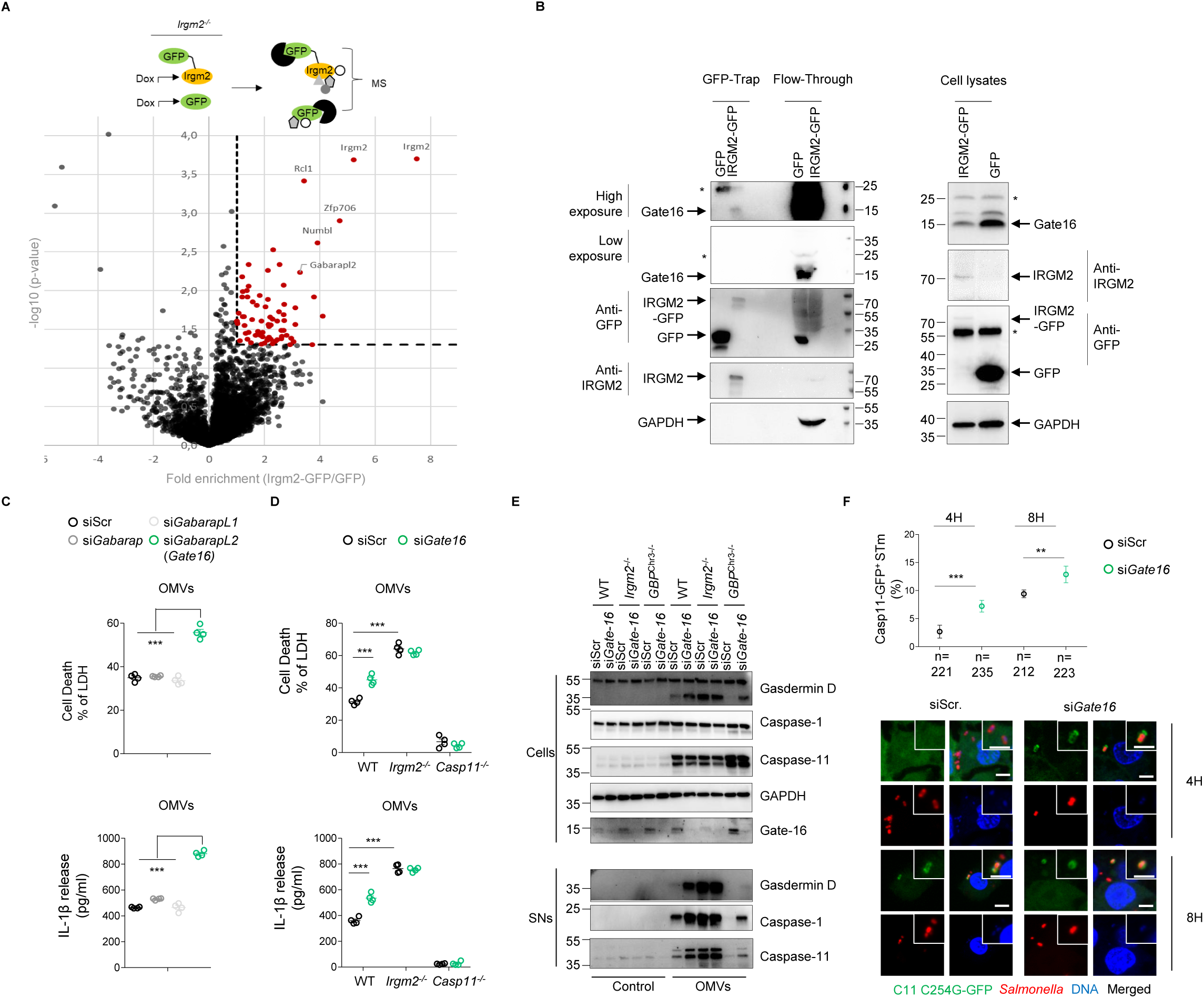
Irgm2 cooperates with Gate-16 to dampen the non-canonical inflammasome activation. **(A)** GFP-Trap coupled to mass spectrometry strategy used. The volcano plot represents 3 independent combined experiments. Threshold selection of enriched proteins in Irgm2-GFP fraction was set at 2 fold enrichment (x axis) and p-value <0.05 (y axis). **(B)** GFP-Trap assay of the presence of Gate16 in Irgm2-GFP-enriched fraction from the lysates of IFNγ-primed i*Irgm2*^−/−^ BMDMs complemented with Irgm2-GFP or GFP. Gate16, Irgm2-GFP and GAPDH are shown in GFP-enriched fractions, flow-through and total cell lysates. **(C)** LDH and IL-1β release from siRNA-treated WT BMDMs, and then exposed to 2.5 µg/2.10^5^ cells of OMVs for 16 hours. **(D)** LDH and IL-1β release from WT, *Irgm2*^−/−^ and *Casp11*^−/−^ BMDMs silenced for *Gate16* and treated for 16 hours with 2,5µg/2.10^5^ cells of OMVs. **(E)** Western blot examination of processed caspase-1 (p20) and gasdermin-D (p30) in supernatants and pro-caspase-1 (p45), pro-gasdermin-D (p55), Gate16 and GAPDH in cell lysates of *Gate16*-silenced WT, *Irgm2*^−/−^ and *GBP*^Chr3–/–^ BMDMs exposed to 2,5.10^5^µg/2.10^5^ cells of OMVs for 16 hours. * Non specific **(F)** Representative confocal fluorescence microscopy images and associated quantifications of caspase-11-C254G-GFP (green) recruitment to *S*.Tm-mCherry (*orgA*^-^, red, MOI 10) in IFNγ-primed iWT BMDMs silenced for *Gate16* after 4 and 8 hours of infection. Nucleus (blue) was stained with Hoescht; scale bar 5µm. For quantifications, the percentage of bacteria positive for caspase-11-C254G-GFP was determined by combining the bacterial counts from n=3 independent experiments and expressed as mean ± SEM. ***p ≤ 0.001 for the indicated comparisons using t-test with bonferroni correction. Data information: Data shown as means ± SEM (Graphs C, D) from n=4 independent pooled experiments; ***p ≤ 0.001 for the indicated comparisons using t-test with bonferroni correction. Image (B) is representative of one experiment performed two times and (E) represents one experiment out of three. (A) represents one experiment out of three independent experiments.

### Gate-16 inhibits non-canonical inflammasome activation in human macrophages

Gate-16 is expressed in both human and rodents, yet human only express one IRGM, although mice have three (Irgm1-3). Therefore, we performed siRNA-based experiments in primary human monocyte-derived macrophages (hMDMs) to determine whether IRGM and GATE16 might also regulate the caspase-4/5 non-canonical inflammasome. Although the use of the caspase-4/5 inhibitor LEVD and of the NLRP3 inhibitor MCC950 showed that hMDMs responded to *Salmonella* infection by activating the non-canonical inflammasome, we failed to observe any regulatory role for IRGM at regulating such process (**Fig 4A, EV 4A**). However, *GATE16* silencing increased their ability to respond to *Salmonella* through the non-canonical inflammasome (**Fig 4A, EV 4A**). When we used Nigericin to trigger the canonical NLRP3 inflammasome, hMDMs knocked down for *GATE16* did not show cell death and IL-1B alterations (**Fig 4B**). To the contrary, *IRGM*-silenced hMDMs had higher IL-1B and cell death levels than their respective controls, which is reminiscent of previous studies that showed a regulatory role for IRGM on the canonical NLRP3 inflammasome (**Fig 4B**). Although, the research of a protein with a similar function of rodent Irgm2 in human warrants further investigations, our results suggest that GATE16 function is conserved between both species. Altogether, our results identified two novel regulators, namely Irgm2 and GATE16 that cooperatively restrict the non-canonical inflammasome in both mice and humans. This suggests that the Irgm2/Gate-16 axis is part of a novel regulatory path that fine-tunes the non-canonical inflammasome threshold level.

**Figure 4:**
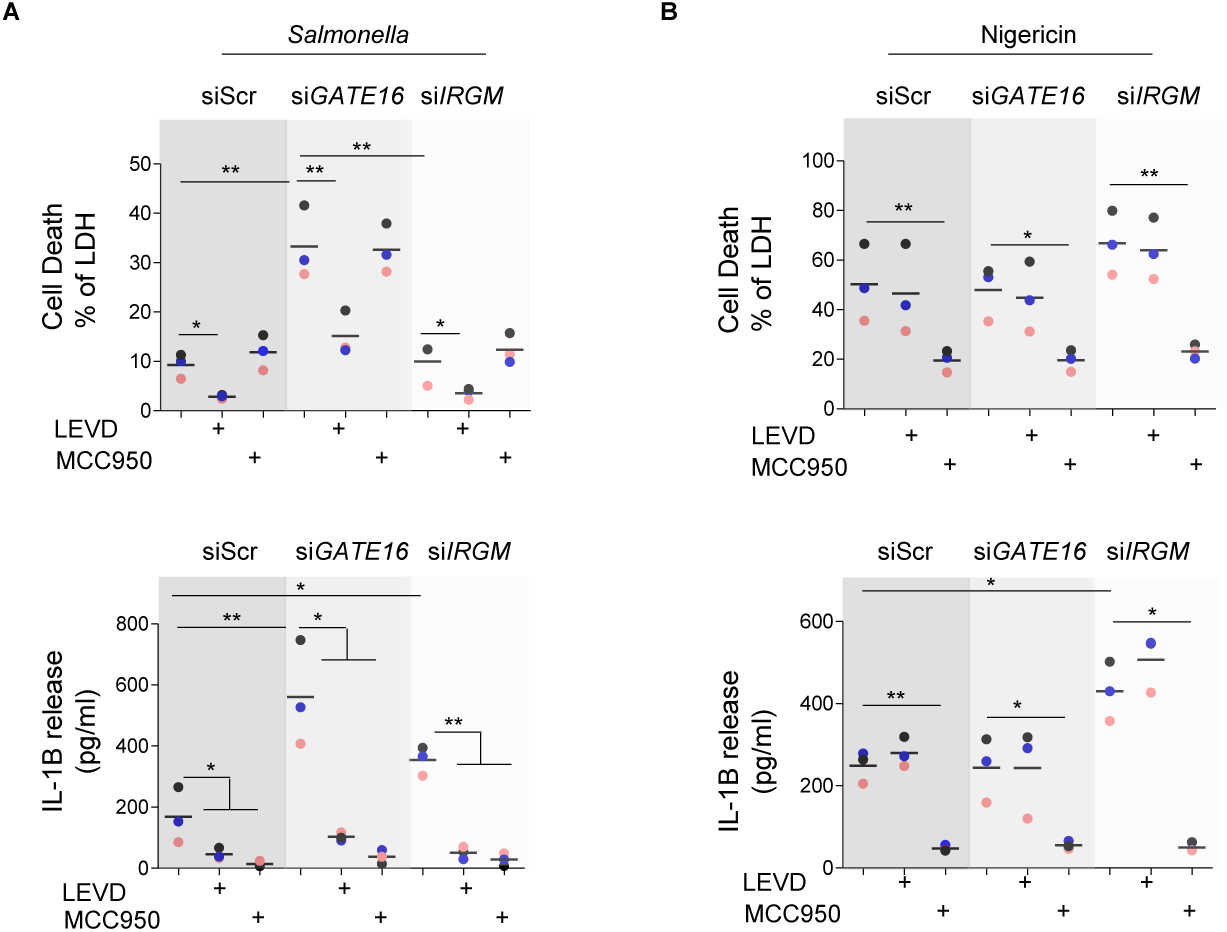
GATE16 inhibits non-canonical inflammasome activation in human macrophages. **(A)** LDH and IL-1β release from siRNA-treated primary human Monocytes-Derived Macrophages (hMDMs) infected with *S*. Typhimurium *orgA*^-^ (MOI25) for 16 hours. When specified, the caspase-4/5 inhibitor Z-LEVD (25µM) or the NLRP3 inhibitor MCC950 (10µM) were added to the experiments. Results from n=3 donors were plotted. **(B)** LDH and IL-1β release from siRNA-treated primary human Monocytes-Derived Macrophages (hMDMs), primed with IFNγ (10UI/mL) and PAM3CSK4 (100ng/mL) and then stimulated with Nigericin (20µM) for 4 hours. When specified, the caspase-4/5 inhibitor Z-LEVD (25µM) or the NLRP3 inhibitor MCC950 (10µM) were added to the experiments. Results from n=3 donors were plotted. Data information: Data shown as means ± SEM (Graphs A, B) from n=3 independent donors; each donor is represented with a coloured circle; * p ≤ 0.05, **p ≤ 0.01, ***p ≤ 0.001 for the indicated comparisons using one-way ANOVA with multiple Bonferroni correction.

## Discussion

Activation of the non-canonical inflammasome is of major importance at triggering an inflammatory response upon gram-negative bacterial challenge. However, uncontrolled non-canonical inflammasome response drives endotoxic shock, which suggests that host factors might tightly regulate the non-canonical inflammasome at various steps. Conversely, two recent studies have uncovered that the IRF2 transcription factor (and to a lower extend IRF1) transcriptionally control murine gasdermin D and human caspase-4 expression (Kayagaki *et al*, 2019; Benaoudia *et al*, 2019). In addition, SERPINB1 has also been found to directly interact and inhibit activation of the inflammatory caspases-1, -4, and -11 (Choi *et al*, 2019). Here we report a critical role of Irgm2 and Gate-16 at balancing the non-canonical inflammasome activity. The Irgm2/Gate-16 axis was required to inhibit caspase-11 recruitment to the gram-negative bacterial product LPS in the host cell cytosol, which provided controlled caspase-11 response and protection against sepsis. While this manuscript was submitted, the group of J Coers also described the regulation of the caspase-11 non-canonical inflammasome by the Irgm2 axis. These findings open many yet unanswered questions such as at which step the Irgm2/Gate16 axis is regulating caspase-11 recruitment to bacterial products. Gate-16 has been found to control proper cytosolic localization of various GBPs (Park *et al*, 2016; Sasai *et al*, 2017), including GBP2, crucial at regulating caspase-11 recruitment on intracellular pathogen PAMPs. However, our results indicate that *Gate16*/*Irgm2* removal in GBP-deficient macrophages reinsures caspase-11-dependent response, which suggests that the Gate-16/Irgm2 path might regulate caspase-11, at least to certain extend, independently of the GBPs. So far, due to the lack of information, a possible function of Irgm2 remains elusive, but Irgm1 and its human homologous IRGM have been described to participate in the autophagy/xenophagy processes (Azzam *et al*, 2017; Maric-Biresev *et al*, 2016). In addition, Gate-16 belongs to the ATG8-like proteins, including LC3 (abc), Gabarap and GabarapL1, all involved in various autophagy/membrane remodelling step regulation such as lysosome biogenesis, autophagosome formation and closure (Nguyen *et al*, 2016; Gu *et al*, 2019; Lee & Lee, 2016). To this regard, both Irgm1/IRGM and Gabarap proteins inhibit activation of the canonical Nlrp3 inflammasome (pei *et al*, 2017; Mehto *et al*, 2019b, 2019a). Therefore, one can hypothesizes that *Gate-16* and *Irgm2* deficiencies could lead to defective autophagy, which would promote cytosolic LPS accumulation and an exacerbated caspase-11 activation. Therefore, experiments using various genetically invalidated cells for autophagy components such as ATG4b, responsible of ATG8 family protein maturations status, might help at determining the importance of the autophagy machinery on Irgm2/Gate-16 axis-inhibited caspase-11 response. Another possible explanation relies on the Golgi enrichment of both Irgm2 and Gate-16 (Sagiv, 2000; Zhao *et al*, 2010). Indeed, Gate-16 also regulates Snare-dependent vesicular trafficking, independently of its autophagy function (Sagiv, 2000). Various groups previously found that endocytosed and intracellular monomeric LPS could be targeted to the Golgi apparatus (Latz *et al*, 2002; Thieblemont & Wright, 1999). Caspase-4, 5 and 11 need accessible lipid A to oligomerize and auto activate, which can be extremely difficult in presence of multimeric and hydrophobic LPS particles to the contrary of monomeric LPS that might present a more accessible lipid A. An attractive hypothesis is that GBP-mediated bacterial membrane damages allow LPS retrieval from aggregates in order to ensure proper lipid A exposure to caspase-11 (Santos *et al*, 2018). Therefore, one could speculate that Golgi-regulated monomeric LPS trafficking might be impaired in absence of either Irgm2 or Gate-16, which would allow direct caspase-11/lipid A interactions without the need for GBPs.

Another standing question lies on the hierarchical involvement of Irgm2 and Gate-16 at regulating the non-canonical inflammasome response. Although our results suggest that both proteins are part of the same pathway, their respective contribution at inhibiting LPS-induced caspase-11 activation remains elusive. Due to the strong response of *Irgm2*-silenced BMDMs compared to *Gate-16* knock down, it is tempting to put Irgm2 as an upstream regulator of Gate-16, which suggests that additional partners/effectors might be involved downstream of Irgm2. In this context, gate16 pull down coupled to mass spectrometry experiments might help identifying additional regulators of the non-canonical inflammasome response.

Our results showed that both murine and human Gate-16 regulate the non-canonical inflammasome response to LPS-containing particles. Yet, we failed to isolate IRGM as a human functional homologous of Irgm2. Given the strong role of Irgm2 at regulating the non-canonical inflammasome in mice, there is a possibility that another, yet unidentified, human protein holds a similar function of the one carried out by Irgm2. Therefore, this warrants future investigations to identify such Irgm2-like human protein.

Human and mice have different sensitivities to LPS. Indeed, LPS-induced sepsis in mice requires 1–25 mg/kg of LPS whereas humans have a 100-1 000 000 time lower sepsis threshold (2–4 ng/kg of LPS) (Fink, 2014). Another explanation could be that the evolutionary loss of Irgm2 in humans would leave human cells with only Gate-16, which would greatly lower the sensitivity of human cells to cytosolic LPS-activated non-canonical inflammasome.

In summary, our work identified two negative regulators of caspase-11 recruitment to bacterial membranes, namely Irgm2 and Gate-16. Additional investigations will be necessary to understand how both effectors balance caspases-4, -5 and -11 sensitivity to intracellular LPS, and what specific physiological and cellular processes Irgm2 and Gate-16 cover together.

## Method

Reagents, biological samples and their concentration of use are referenced in the Appendix table 1

### Mice

*Casp11*^−/−^, *Casp1*^−/−^*Casp11*^−/−^, *Nlrp3*^−/−^, and *GBP*^Chr3–/–^ mice have been described in previous (Yamamoto *et al*, 2012; Martinon *et al*, 2006; Li *et al*, 1995; Wang *et al*, 1998). *Irgm2*^−/−^ mice provided from the Jackson laboratory (USA). All mice were bred at the IPBS institute (Toulouse, France) animal facilities according to the EU and French directives on animal welfare (Directive 2010/63/EU). Charles Rivers provided WT C57B6 mice.

### Animal sepsis models

8-12 weeks old mice (sex-matched, 6-10 per group) were injected intraperitoneally with a solution of 100µL of poly(IC) LMW (invivogen, 100µg/animal) for 6 hours. Then, 5mg/kg of LPS (Invivogen, O111:B4) or 5 or 25µg/animal of outer membrane vesicles (*E. coli*, Invivogen) were intraperitoneally injected. Mice survival was monitored over 80 hours. For cytokine assays, poly(IC)-primed mice were injected with 25µg/animal of OMVs for 8 hours and plasma cytokines amounts were addressed using ELISA kits (listed in the Appendix Table 1). There were no randomization or blinding performed.

Animal experiments were approved (License APAFIS#8521-2017041008135771, Minister of Research, France) and performed according to local guidelines (French ethical laws) and the European Union animal protection directive (Directive 2010/63/EU).

### Animal infection

6-10 mice/group were intraperitoneally infected with 5.10^5^ CFUs of stationary phase *Salmonella sifA*^-^ strain. Animals were sacrificed 48 hours after infection. Spleen bacterial loads were evaluated using CFU plating and plasma cytokine levels were determined using ELISA kits. There were no randomization or blinding performed.

### BMDM isolation and culture

Murine Bone Marrow-Derived Macrophages (BMDMs) generation has previously been described. Briefly, bone marrow progenitors were differentiated in DMEM (Invitrogen) supplemented with 10% v/v FCS (Thermo Fisher Scientific), 10% v/v MCSF (L929 cell supernatant), 10 mM HEPES (Invitrogen), and nonessential amino acids (Invitrogen) for 7 days. For experiments, 1.25×10^6^, 2.5×10^5^, or 5×10^4^ BMDMs were seeded in 6-, 24-, or 96-well-plates, respectively. When described BMDMs were pre-stimulated overnight with either PAM3CSK4 (InvivoGen, 100 ng/mL) or IFN-ɣ (PeProtech, 100UI/mL). For non-canonical inflammasome stimulation, we used pure LPS (O111B4, Invivogen, 1µg/mL), Outer Membrane Vesicles (*E. coli*, Invivogen, 0.5, 1 and 2.5 µg/2.10^5^ cells) or various gram-negative bacterial strains were used including, *Shigella flexneri* (M90T, MOI25), *Salmonella* Typhimurium *orgA*^-^ (SL1344, MOI25), *E. coli* (K12, MOI25), *C. koseri* (MOI25), *E. cloacae* (MOI25).

For canonical inflammasome stimulations, over-night (ON) IFNγ-primed BMDMs were then prestimulated with PAM3CSK4 (InvivoGen, 100 ng/mL) for 4hours to induce pro-IL1β expression. Then, Nigericin (NLRP3 activator, 40µM, Invivogen), dA:dT (AIM2 activator, 1µg/mL, Invivogen), TcdB toxin (PYRIN inducer, 0.05µg/mL, List Biological Laboratories) or flagellin (NLRC4 trigger, 1µg/mL, Invivogen) were used to stimulate various canonical inflammasomes. Both flagellin and poly (dA:dT) were transfected into cells using FuGeneHD (Promega) transfection reagent in Opti-MEM culture medium. When specified *P. aeruginosa* (PAO1) and *S*. Tm strains (MOI 5) were used to trigger NLRC4 inflammasome response.

For all stimulations, macrophages medium was replaced by serum-free and antibiotic-free Opti-MEM medium and inflammasome triggers were added to the macrophages for various times.

Specific to infections, plates were centrifuged for 1 minute, 800 rpm to ensure homogenous infections. Then, extracellular bacteria were eliminated with gentamicin (100 µg/ml, Invitrogen).

### Bacterial cultures

bacteria were grown overnight in Luria Broth (LB) medium at 37°C with aeration and constant agitation in the presence or absence of antibiotics (specified in the Appendix table 1). Stationary phase (OD of 2-2.5) bacteria when then used for infections. Stimulation of the NLRC4 inflammasome by *S*. Typhimurium SL1344 and *P. aeruginosa* PAO1 bacteria required proper T3SS and flagellin expression; therefore bacteria were sub-cultured the next day by dilution overnight culture 1/50 and grew until reaching an O.D600 of 0.6 - 1

### Gene knock down

Gene silencing was achieved using siRNA pools (Dharmacon, 25nM/well listed in table S1) as previously described (Santos *et al*, 2018; Meunier *et al*, 2015) or accell siRNA technology. SiRNA smart pools from Dharmacon were transfected into cells using the DharmaFECT 4 transfection reagent (Dharmacon) for 48 hours. Primary Human macrophages were treated with 1µM siRNA Accell (Dharmacon, smart pool) in absence of transfection reagent for 72 hours. Then, murine BMDMs and human macrophages were stimulated with 1µg/2.10^5^ cells of OMVs or infected with *Salmonella* Typhimurium (*orgA*^-^) to trigger non-canonical inflammasome response. For siRNA experiments, gene knockdown efficiency was monitored by qRTPCR or immunoblotting (WB) assays.

### Quantitative Real Time PCR

Cellular RNA were extracted from 2.5 10^5^ cells using RNeasy Mini Kit (Qiagen). mRNAs were reverse transcribed with the Verso cDNA Synthesis Kit (ThermoScientific). Regarding qPCR experiments, 1µM of primers (Appendix Table S1), SYBR™ Select Master Mix (ThermoScientific) and 15ng of cDNA were mixed in a 10µL reaction in a QuantStudio 5 device (Applied Biosystems). Primers were generated using primer3 software.

### Cytokine and pyroptosis measurement

Murine Il-1α, IL-1β, TNFα, IL12, IL18, IFNγ, IL-6 and human IL-1B cytokine levels were measured by ELISA (listed in appendix table 1). LDH Cytotoxicity Detection Kit (Takara) allowed to monitore for cell lysis. Normalization of spontaneous lysis was calculated as follows: (LDH infected – LDH uninfected)/(LDH total lysis – LDH uninfected)*100.

### Immunoblotting

Preparation of cell lysates and supernatants have been described previously. Proteins were loaded and separated in 12% SDS-PAGE gels and then transferred on PVDF membranes. After 1 hour of saturation in Tris-buffered saline (TBS) with 0.05% Tween 20 containing 5% non-fat milk (pH8), membranes were incubated overnight with various antibodies (referenced in table 1). The next day, membranes were washed 3 times in TBS 0.1% Tween 20 and incubated with appropriate secondary horseradish peroxidase (HRP)-conjugated antibody (dilution 1/5000-10000, listed in Appendix table 1) for 1h at room temperature. Then, after 3 washes, immunoblottings were revealed with a chemiluminescent substrate ECL substrate (Biorad) and images were acquired using ChemiDoc Imaging System (Biorad). All antibody references and working dilutions are presented in Appendix table 1.

### Microscopy

2.5×10^5^ BMDMs on glass coverslips were infected with *S*. Typhimurium (MOI10) expressing an mCherry fluorescent protein. At the indicated times, cells were washed three times with PBS and fixed with 4% PFA for 10 minutes at 37°C. 0,1M Glycine was used to quench excess of PFA for 10 min at room temperature. Then cells were permeabilized and incubated with primary antibodies O/N at 4°C in Saponin 0.1% /BSA 3% solution. Cellular stainings were achieved using Hoescht (DNA labeling), GBP2 antibody (gift from J Howard). Coverslips were then washed with Saponin/BSA solution and further incubated with the appropriate secondary antibodies coupled to fluorochromes (1/1000, appendix table 1). After 3 washes with PBS, cells were mounted on glass slides using Vectashield (Vectalabs). Coverslips were imaged using confocal Zeiss LSM 710 (Image core Facility, IPBS, Toulouse or an Olympus/Andor CSU-X1 Spinning disk microscope using a 63x oil objective. Otherwise specified, 5-10 fields /experiment were manually counted using Image J software.

### Transduction of iBMDMs

HEK 293-based retroviral packaging cell line (GP2-293) were plated in 10 cm petri dish in DMEM + 10% FCS + 1 % PS. When cell’s confluency reached 60-80%, cells were placed in serum and antibiotics-free Opti-mem medium and transfected with VSV-G encoding vector (pMD.2G) along with CASP11-C254G-GFP or pRetro (-GFP or – Irgm2-GFP) vectors using PEI transfection reagent. 10 hrs after transfection cell medium was replaced by DMEM + 10% FCS + 1 % PS. At 48 hrs post-transfection cell’s supernatant containing retroviral particles were collected, filtered 0.45µm and used to transduce target cells. After 48 hours, puromycin (5µg/mL) was used to select cells positively transduced with the transgene. When vectors contained GFP fusions, cells were sorted using Fluorescence-Activated Cell Sorting.

### Immunoprecipitation and GFP Trap

*Irgm2*^−/−^ immortalized macrophages were transduced with retroviral vectors carrying a doxycycline inducible Irgm2-GFP, or GFP alone constructs, cloned into Retro-X™ Tet-On® 3G vector (Clontech Laboratories, Inc.). To ensure proper Irgm2-GFP expression cells were incubated 16 hrs with doxycycline 1µg/ml in presence of IFNɣ. Irgm2-GFP and associated protein complexes were pull-down using GFP-Trap magnetic beads according to manufacturer’s instructions (chromotek). Briefly, cells were lysed in CoIP lysis buffer (10 mM Tris/Cl pH7.5; 150 mM NaCl; 0.5mM EDTA; 0.5% NP40, 0.09% Na-Azide) supplemented with a protease inhibitor cocktail (Roche). Cell lysates were then incubated with GFP-Trap-MA beads for 1 hr at 4°C. After 2 washes with wash-buffer (10 mM Tris/Cl pH 7.5; 150 mM NaCl; 0.5 mM EDTA, 0.018% Na-Azide) GFP-Trap complexes were boiled for 10 min at 95°C in RIPA buffer + laemli before separation on SDS-PAGE and mass spectrometry or immunoblotting.

### Mass spectrometry analysis

Immunopurified protein samples were reduced with β-Mercaptoethanol by heating at 95°C for 5min, and cysteines were alkylated by addition of 90mM iodoacetamide. Samples were loaded on a 1D SDS-PAGE gel and proteins were isolated in a single gel band, which was excised and washed with several cycles of 50 mM ammonium bicarbonate-acetonitrile (1:1). Proteins were in-gel digested using 0.6 µg of modified sequencing grade trypsin (Promega) in 50 mM ammonium bicarbonate overnight at 37°C. Resulting peptides were extracted from the gel by successive incubations in 50 mM ammonium bicarbonate and 10% formic acid-acetonitrile (1:1), then dried in a speed-vac, and resuspended with 22µL of 5% acetonitrile, 0.05% trifluoroacetic acid (TFA) for MS analysis. Peptides were analyzed by nanoLC-MS/MS using an UltiMate Nano/Cap System NCS-3500RS coupled to a Q-Exactive HFX mass spectrometer (Thermo Fisher Scientific, Bremen, Germany). Separation was performed on a C-18 column (75 µm ID x 50 cm, Reprosil C18) equilibrated in 95% solvent A (5% acetonitrile, 0.2% formic acid) and 5% solvent B (80% acetonitrile, 0.2% formic acid), using a gradient from 10 to 45% gradient of solvent B over 60 min at a flow rate of 350 nL/min. The mass spectrometer was operated in data-dependent acquisition mode with the Xcalibur software. Survey MS scans were acquired in the Orbitrap on the 350– 1400 m/z range, with the resolution set to 60,000, and the 12 most intense ions were selected for fragmentation by Higher-energy Collisional Dissociation (HCD) using a normalized collision energy of 28. MS/MS scans were collected at 15,000 resolution with an AGC target value of 1e5 and a maximum injection time of 22 ms. Dynamic exclusion was used within 30 s to prevent repetitive selection of the same peptide. Three replicate MS analysis were performed for each sample.

### Bioinformatic processing of mass spectrometry data

Raw mass spectrometry files were searched using Mascot (Matrix Science) against the Mouse entries of the Swissprot-TrEmbl protein database. The enzyme specificity was “trypsin”, with a maximum of 2 misscleavages. Cysteine carbamidomethylation was set as a fixed modification, and N-terminal protein acetylation and methionine oxidation were specified as variable modifications. For the search, mass tolerance parameters were set at 5 ppm on the parent ion and 20 mmu on the fragment ions. Protein identification results were then validated with the Proline software by the target-decoy approach using a reverse database at a both a peptide and protein FDR of 1%. To perform label-free relative quantification of proteins, the “abudance” metric retrieved by Proline was used, after global normalization of the MS signal across all MS runs. For each protein, a mean abundance value was computed from technical LC-MS replicate runs, and log2-transformed. Missing protein abundance values were then replaced by a noise value estimated for each analysis as the 1% lowest percentile of the protein abundance values distribution. Bona fide Irgm2 interactors were identified by comparing Irmg2-GFP immuno-purified samples and GFP control samples. For each protein, an enrichment ratio relative to the control and a Student t-test p-value were calculated from the protein abundance values derived from 3 independent biological replicate experiments. Relevant interactors were selected based on an enrichment ratio higher than 2 and a Student t-test p-value lower than 0.05.

### Genetic invalidation of *Caspase11* and *Irgm2* genes in immortalized BMDMs

*Casp11* and *Irgm2* genes were knocked-out using the crispr/cas9 system in onco J2-immortalized (i) bone-marrow derived macrophages (BMDMs) iWTs or i*Irgm2*^−/−^ macrophages. Single guide RNAs (sgRNA) specifically targeting Caspase-11 exon 2 forward (5’CACCGCTTAAGGTGTTGGAACAGCT3’) reverse (5’AAACAGCTGTTCCAACACCTTAAGC3’), Irgm2 exon 2 forward (5’CACCGTTCCATGTTGTCGAGCAACG3’) reverse (5’AAACCGTTGCTCGACAACATGGAAC3’) were designed using Benchling tool (Benchling.com), and oligonucleotides were synthesized by Sigma-Aldrich. Crispr guide RNA oligonucleotides were then hybridized and cloned in Lenti-gRNA-Puromycin vector using BsmBI restriction enzyme (lentiGuide-Puro, Addgene 52963, Feng Zhang lab). HEK293T cells were transfected for 48 hours with all constructs (lipofectamine 2000) together with the lentiviral packaging vector p8.91 (Didier Trono lab, EPFL, Switzerland) and the envelop coding VSVg plasmid (pMD.2G, Addgene 12259, Didier Trono lab). 48 hours later, viral supernatants were harvested and subsequently filtered on 0.45 µm filter. Recipient cells expressing Cas9 (1,000,000 cells/well in 6-well plates) were generated using lentiviral transduction with a Cas9-expressing lentiviral vector (lentiCas9-Blast, Addgene 52962, Feng Zhang lab). The, Cas9^+^ cells were infected with packaged viral particles. To ensure efficient infection viral particles were centrifugated for 2 h at 2900 rpm at 32°C in presence of 8μg/ml polybrene. 48 h later, medium was replaced and Puromycin selection (10µg/mL) was applied to select positive clones for two weeks. Puromycin-resistant cells were sorted at the single cell level by FACS (Aria cell sorter). Individual clones were subjected to western blotting to confirm the absence of targeted proteins.

### Generation of human monocyte-derived Macrophages

Peripheral Blood Mononuclear Cells (PBMCs) were isolated from buffy coat of healthy donors obtained from the EFS Toulouse Purpan (France) as described previously (ref). Briefly, PBMCs were isolated by centrifugation using standard Ficoll-Paque density (GE Healthcare). The blood was diluted 1:1 in phosphate-buffered saline (PBS) pre-warmed to 37°C and carefully layered over the Ficoll-Paque gradient. The tubes were centrifuged for 25 min at 2000 rpm, at 20°C. The cell interface layer was harvested carefully, and the cells were washed twice in PBS (for 10 min at 1200 rpm followed by 10 min at 800 rpm) and re-suspended in RPMI-1640 supplemented with 10% of foetal calf serum (FCS), 1% penicillin (100 IU/mL) and streptomycin (100 µg/ml). Monocytes were separated from lymphocytes by positive selection using CD14+ isolation kit (Myltenyi biotec). To allow differentiation into monocyte-derived macrophages, cells were cultured in RPMI medium (GIBCO) supplemented with 10% FCS (Invitrogen), 100 IU/ml penicillin, 100µg/ml streptomycin, 10 ng/ml M-CSF for 7 days.

### Ethics statements

The use of human cells was approved by the Research Ethical Committee, Haute-Garonne, France. Buffy coats were provided anonymously by the EFS (établissement français du sang, Toulouse, France). Written informed consent was obtained from each donor under EFS contract n° 21PLER2017-0035AV02, according, to “Decret N° 2007-1220 (articles L1243-4, R1243-61)”.

### Statistical analysis

Statistical data analysis was performed using Prism 5.0a (GraphPad Software, Inc.). T-test with Bonferroni correction was used for comparison of two groups. For multiple comparisons, One-Way ANOVA with Multiple Bonferroni correction test was used. Data are reported as mean with SEM. For animal experiments Mann-Whitney tests were performed and for mouse survival analysis log-rank Cox-Mantel test was selected. P values are given in figures, NS means non-significant. Significance is specified as *p ≤ 0.05; **p ≤ 0.01, ***p ≤ 0.001.

## ACKNOWLEDGEMENTS

We would like to acknowledge Jonathan Howard for anti-Irgm2 anti-serum and anti-GBP2 as well as Biotem company for generating anti-Irgm2 antibodies; Junying Yuan (Harvard Med School, Boston, USA) and B. Py (CIRI institute, Lyon, France) for *Caspase1*^−/−^*Caspase11*^−/−^ and *Caspase11*^−/−^ mice (Li *et al*, 1995; Wang *et al*, 1998); M. Yamamoto (Osaka University, Japan) for *GBP*^Chr3–/–^ mice (Yamamoto *et al*, 2012), V. Petrelli and B. Py for the *Nlrp3*^−/−^ mice (Martinon *et al*, 2006). *Irgm2*^−/−^ mice came from the Jackson laboratory. Authors also acknowledge the animal facility, mass spectrometry and microscopy platforms of the IPBS institute. We specifically acknowledge Drs. C. Cougoule and Y. Rombouts for fruitful discussions and suggestions as well as for reading and implementing the MS. This project was funded by grants from FRM “Amorçage Jeunes Equipes” (AJE20151034460), ERC StG (INFLAME 804249) and ATIP to EM and from the European Society of Clinical Microbiology and Infectious Diseases (ESCMID, 2020) to RP. MY (Masahiro Yamamoto) is supported by the Research Program on Emerging and Re-emerging Infectious Diseases (JP19fk0108047), Japanese Initiative for Progress of Research on Infectious Diseases for global Epidemic (JP19fm0208018) and Strategic International Collaborative Research Program (19jm0210067h) from Agency for Medical Research and Development (AMED), Grant-in-Aid for Scientific Research on Innovative Areas (Production, function and structure of neo-self; 19H04809), for Scientific Research (B) (18KK0226 and 18H02642) and for Scientific Research (A) (19H00970) from Ministry of Education, Culture, Sports, Science and Technology of Japan.

## AUTHOR CONTRIBUTIONS

EE and EM designed the experiments with the help of RP. EE, RP and EM wrote the manuscript. EE and RP performed the experiments with the help of SB, PJB, AH, KS and MP. KC and OBS performed essential mass spectrometry run acquisitions and analysis. MY and JCH provided essential reagents to conduct the project.

## Figures

**Expanded Views (EV):**

**EV1: Irgm2 specifically controls non-canonical inflammasome response**

**(A)** qRT-PCR measurement of silencing efficacy on the mRNA levels of *Irgm1-3, Gbp2* and *Caspase-1* in BMDMs, prestimulated with 100UI/ml of IFNγ for 16 hours. N=3 independent experiments normalized to β-actin mRNA levels. Data are expressed as mean ± SEM.

**(B)** Immunoblotting of Irgm2, Caspase-1, Caspase-11 and GAPDH expression in IFNγ-primed WT or *Irgm2*^−/−^ BMDMs. Image represents one experiment performed two times. * Non specific

**(C)** Immunoblotting of GBP2, GBP5 and GAPDH expression in LPS- or IFNγ-primed WT or *Irgm2*^−/−^ BMDMs. Image represents one experiment performed two times. * Non specific

**(D)** LDH and IL-1β release from WT, *Irgm2*^−/−^, *Casp11*^−/−^ *and Nlrp3*^−/−^ BMDMs treated for 16 hours with 2,5µg/2.10^5^ cells of OMVs in presence or not of 10µM of MCC950 (NLRP3 inhibitor). Data are expressed as mean ± SEM from n = 4 experiments.

**(E)** Immunoblots of *Casp11* or *Irgm2* deletion efficacy in immortalized BMDMs. Image represents one experiment performed two times. * Non specific

**(F)** Release of LDH and IL-1β from IFNγ- and PAM3CSK4-primed WT, *Irgm2*^−/−^, *Casp11*^−/−^ or *Casp1*^−/−^*Casp11*^−/−^ BMDMs transfected (using FuGeneHD) with flagellin or poly(dA:dT) or stimulated with either Nigericin or TcdB toxin for 6 hours. Data are expressed as mean ± SEM from n = 4 experiments.

**(G)** Western blot examination of processed caspase-1 (p20) and gasdermin-D (p30) in supernatants and pro-caspase-1 (p45), pro-gasdermin-D (p55) and GAPDH in cell lysates of WT and *Irgm2*^−/−^ BMDMs infected for 4 h with *S*. Typhimurium and *P. aeruginosa* (NLRC4 inflammasome). Image represents one experiment performed two times. * Non specific

**(H)** Cytokine levels in plasma from WT, *Casp11*^−/−^ and *Irgm2*^−/−^ (n=6 mice per condition) primed with 100µg poly(I:C) for 6 hours and injected i.p with 25 µg of OMVs for 5 hours. Graphs represent one experiment out of three independent experiments; *p ≤ 0.05; **p ≤ 0.01, ***p ≤ 0.001, Mann-Whitney analysis test.

**EV2: *Irgm2* deficiency drives GBP-independent caspase-11 targeting to gram-negative bacterial membranes.**

**(A)** Immunoblots of *Irgm2* silencing efficacy in primary BMDMs. Image represents one experiment performed two times.

**(B)** Cell death (LDH) and IL-1β release evaluation in WT, *GBP*^Chr3–/–^, *Casp11*^−/−^ and *Casp1*^−/−^*Casp11*^−/−^ BMDMs infected for 16 hours with either *S*.Tm *orgA*^-^ or *F. novicida* (MOI 25). Data are expressed as mean ± SEM from n = 4 experiments.

**(C)** Immunoblots of *Irgm2* deletion efficacy in immortalized *GBP*^Chr3–/–^ BMDMs. Image represents one experiment performed two times. * Non specific

**(D)** Kinetic of *S*. Tm (*orgA*^-^)-induced cell death (% LDH release) in IFNγ-primed iWT, i*Irgm2*^−/−^, i*GBP*^Chr3–/–^ and i*GBP*^Chr3–/–^*Irgm2*^−/−^ BMDMs. Data represent one experiment out of n=3 experiments and is expressed as mean ± SEM

**(E)** Confocal fluorescence microscopy images and associated quantifications of caspase-11-C254G-GFP (green) recruitment to *S*.Tm-mCherry (*orgA*^-^, red) in IFNγ-primed iWT, i*Irgm2*^−/−^ i*GBP*^Chr3–/–^ and i*GBP*^Chr3–/–^*Irgm2*^−/−^ BMDMs after 4 hours of infection. Nucleus (blue) was stained with Hoescht; scale bar 5µm. For quantifications, the percentage of bacteria positive for caspase-11-C254G-GFP was determined by combining the bacterial counts from n=3 independent experiments and expressed as mean ± SEM. ***p ≤ 0. for the indicated comparisons using t-test with bonferroni correction.

**EV 3: Gate-16 specifically inhibits the non-canonical inflammasome**

**(A)** Representation of top 11 enriched proteins isolated in Irgm2-GFP fraction. N= 3 independent experiments (Set1-3)

**(B)** qRT-PCR measurement of silencing efficacy on the mRNA levels of *Gabarap, GabarapL1 and L2 (GATE16)* in BMDMs, prestimulated with 100UI/ml of IFNγ for 16 hours. N=3 independent experiments normalized to β-actin mRNA levels. Data are expressed as mean ± SEM.

**(C)** Immunoblots of *GATE16* silencing efficacy in BMDMs. Image represents one experiment performed two times. * Non specific

**(D)** Release of IL-1β from IFNγ (100UI/mL) and PAM3CSK4 (100ng/mL)-primed WT and *Nlrp3*^−/−^ BMDMs treated with 20µM of Nigericin for 4 hours. Data are expressed as mean ± SEM from n = 4 experiments.

**(E)** Representative confocal fluorescence microscopy images and associated quantifications of caspase-11-C254G-GFP (green) recruitment to *S*.Tm-mCherry (*orgA*^-^, red, MOI 10) in IFNγ-primed i*Irgm2*^-/-^ BMDMs silenced for *Gate16* after 4 hours of infection. Nucleus (blue) was stained with Hoescht; scale bar 5µm. For quantifications, the percentage of bacteria positive for caspase-11-C254G-GFP was determined by combining the bacterial counts from n=3 independent experiments and expressed as mean ± SEM. ***p ≤ 0.001 for the indicated comparisons using t-test with bonferroni correction.

**EV 4: Efficacy of h*GATE16* silencing in monocyte-derived macrophages**

**(A**) qRT-PCR measurement of silencing efficacy on the mRNA levels of h*GATE16 and* IRGM in hMDMs, prestimulated with 10UI/ml of IFNγ for 8 hours. N=3 independent experiments normalized to β-actin mRNA levels. Data are expressed as mean ± SEM.

## Graphical abstract

Caspase-11 target cytosolic Gram-negative bacterial LPS, which promotes pyroptosis and IL-1 cytokine maturation and release. IFN-inducible GTPases promote caspase-11 enrichment on bacterial membranes. To the contrary, IFN-inducible Irgm2 and the non-canonical autophagy protein Gate-16 cooperate to refrain unnecessary caspase-11 targeting to Gram-negative bacteria.

